# Allele Frequency Mismatches and Apparent Mismappings in UK Biobank SNP Data

**DOI:** 10.1101/2020.08.03.235150

**Authors:** James Kunert-Graf, Nikita Sakhanenko, David Galas

## Abstract

We report here some anomalies discovered in the minor allele frequencies (MAFs) and some likely mismappings found in our analyses of UK Biobank dataset (UKB) and several other databases. We compared the MAFs present in the UKB to those measured in two other UK studies, ALSPAC and TwinsUK, and found a large set of SNPs for which the UKB MAFs are inconsistent. Additionally, even after accounting for population structure effects and other possible causes of spurious correlations, we found many SNPs that appear to be in interchromosomal linkage. Analyzing these interchromosomal linkages carefully, we found that they are all associated with identical sequences on different chromosomes, implying that these SNPs are simply mismapped. Some (but certainly not all) of the MAF disagreements appear to be the result of these mismappings. Our results, including lists of SNPs with inconsistent MAFs and/or apparent interchromosomal linkage, are freely available to download at: http://kunertgraf.com/data/biobank.html

## 1 Introduction

The UK Biobank dataset, which contains both genotype and phenotype data for approximately 500,000 individuals [1], has an unprecedented scale. This amount of data confers significantly enhanced statistical power that enables novel analyses for garnering biological insights, such as those that seek to characterize the role of complex epistatic interactions in determining traits [2, 3]. The development of novel methods for genetic data analysis, however, requires special care to be taken in both the quality control of the input data and the interpretation of the results. To this end, this study details anomalies within the UK Biobank genetic data, which we initially detected in the process of such methodological development, that we here will characterize in depth.

Our initial project was focused on searches for non-pairwise, multivariable relationships between genotypes and phenotypes within the UK Biobank dataset. The precise nature and goals of this project are not crucial to this study, so we will not detail it further here. This paper focuses instead on some insights into the datasets that we came across in the process of this analysis, and that are more broadly important to questions of quality control, preprocessing and the use of the Biobank data. Specifically, the search for interchromosomal multigenic interactions yielded SNPs that had (i) minor allele frequencies inconsistent with those found by other studies, even studies drawn from similar UK populations; (ii) apparent interchromosomal linkage disequilibrium (ILD), much of which may be explained by mismapping between genes and their related pseudogenes.

This paper identifies 2,323 SNPs with MAFs that are inconsistent with other studies of UK populations, as well as a partially overlapping set of 1,675 SNPs with apparent ILD. Crucially, we find that these SNPs are not detectable by any standard pre-processing or quality control method. We have therefore made our results, including lists of the implicated SNPs, freely available for download at http://kunertgraf.com/data/biobank.html, to enable researchers to easily filter them out for future analyses.

## 2 Minor Allele Frequencies

### 2.1 The MAF mismatch in rs201947198

Our initial study of potential multigenic interactions flagged several SNPs as potentially interesting, including the SNP rs201947198. However, checking the entry for this SNP in the NCBI dbSNP database immediately raised concerns. As shown in Figure 1, dbSNP reports an extremely rare minor allele frequency (MAF) within the ExAC database (MAF=0.00002). This rare a variant should have been filtered out by our preprocessing, but we then confirmed that this SNP had a MAF of 0.375 in our dataset. Figure 1 shows this information as displayed within the Biobank Showcase.

**Figure 1:**
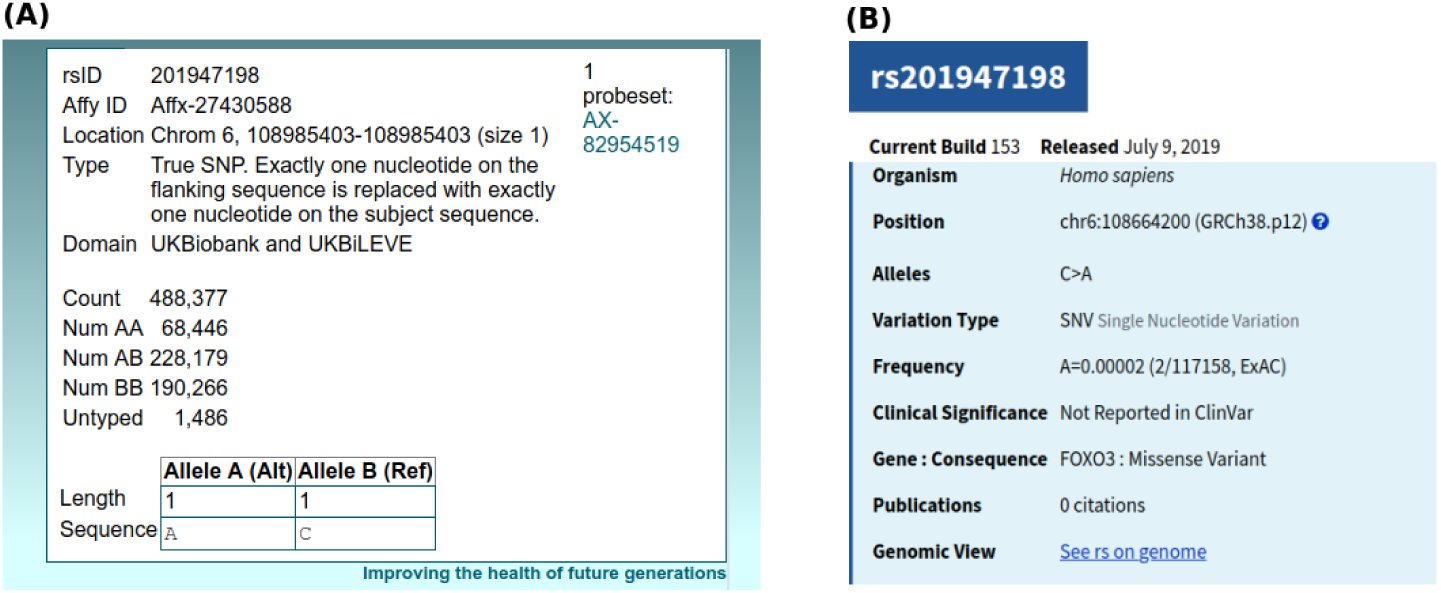
Summary of results for rs201947198: from both **(A)** the UK Biobank, with a MAF=0.375 and **(B)** dbSNP, with a MAF=0.00002.

### 2.2 Comparing MAFs in UKB to several dbSNP cohorts

We suspected that the MAF mismatch found in rs201947198 indicated a potential mismapping so this SNP should be filtered out, and therefore we made an effort to identify and confirm any other SNPs that might be mismapped. Previously, about 300 SNPs were identified by [1] as having an inconsistent MAF when compared to the ExAC database; however, the ExAC cohort is drawn from a different population than the UKB, which might explain some of the expected level of disagreement between the MAFs detected in each study. Fortunately, there exist two cohorts, cataloged by dbSNP [4], that consider populations similar to the UKB: TwinsUK and ALSPAC [5, 6, 7]. We downloaded the full dbSNP database and did a comparison of MAFs among these databases.

Of the 784,256 autosomal SNPs measured by the UK Biobank, 632, 307 (referred to as *N*_*snp*_) were reported in dbSNP for the TwinsUK and ALSPAC cohorts (Note: both cohorts were sequenced by the UK10k project using an Illumina HiSeq platform [7]). The left panel of Figure 2 shows a comparative plot of the MAFs of these SNPs as measured in TwinsUK and ALSPAC (as per dbSNP). These two studies show fairly close agreement, with the width of the distribution quite plausibly due to statistical fluctuations. The right panel, however, compares the TwinsUK database against the UK Biobank and shows a number of clear outliers, for which the MAFs disagree sharply.

**Figure 2:**
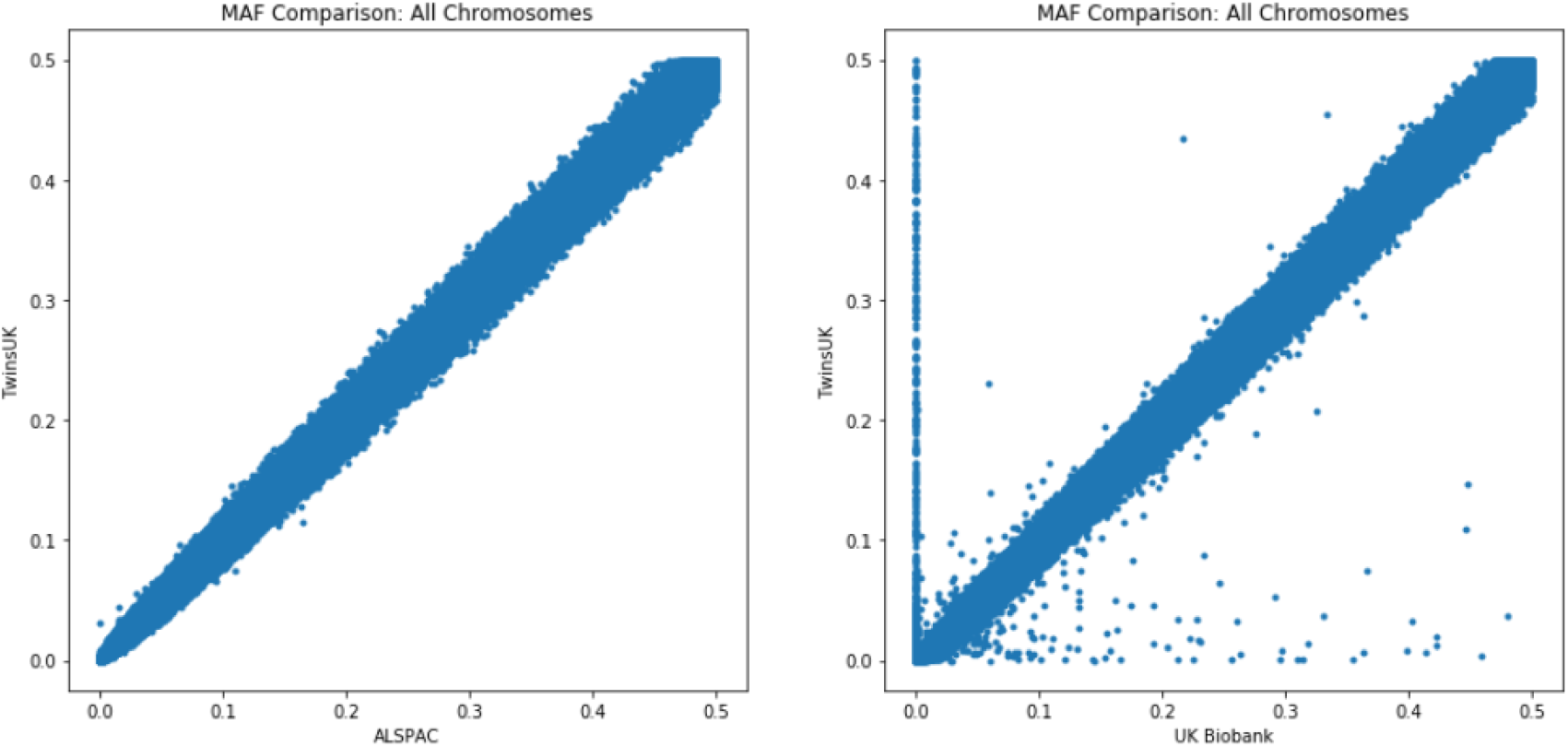
**Left:** MAF values as measured in TwinsUK plotted against those measured in ALSPAC. **Right:** MAF values of TwinsUK plotted against those measured in the UK Biobank.

A higher level of overall noise does not seem like a reasonable explanation for this disagreement, and there is strong evidence of some systematic error. First, there are SNPs that have a MAF of essentially zero in the UK Biobank but are common (much higher MAF) within the TwinsUK and ALSPAC databases. This is notable and calls for an explanation, but in order to avoid disparity in results these SNPs are easily filtered out by simply requiring a MAF threshold in pre-processing. More troublesome is the converse phenomenon: SNPs that are common in the UK Biobank, but very uncommon in the other studies. These SNPs are extremely rare in other studies but have common MAF values in the UK Biobank data. Can this disagreement, and any potentially induced problems, be caught by other pre-processing steps?

To answer this question, we first need a principled method for identifying which SNPs disagree with the MAF values measured in other studies. We do this by making use of the reasonable expectation that the MAFs between two studies should be approximately related by a binomial distribution. Specifically, if we measure MAFs, *p*_*A*_, in study A and *p*_*B*_, in study B, we assume we can take *p*_*A*_ as a parameter of a binomial distribution and use this to calculate the probability of measuring an allele frequency of *p*_*B*_. This assumes random sampling and no other source of disagreement between the two data sets being compared. As shown in Figure 3, the boundaries calculated from this distribution match the shape of the TwinsUK vs. ALSPAC distribution fairly well. We can therefore reasonably use these same boundaries to classify SNPs as outliers in the TwinsUK vs Biobank data.

**Figure 3:**
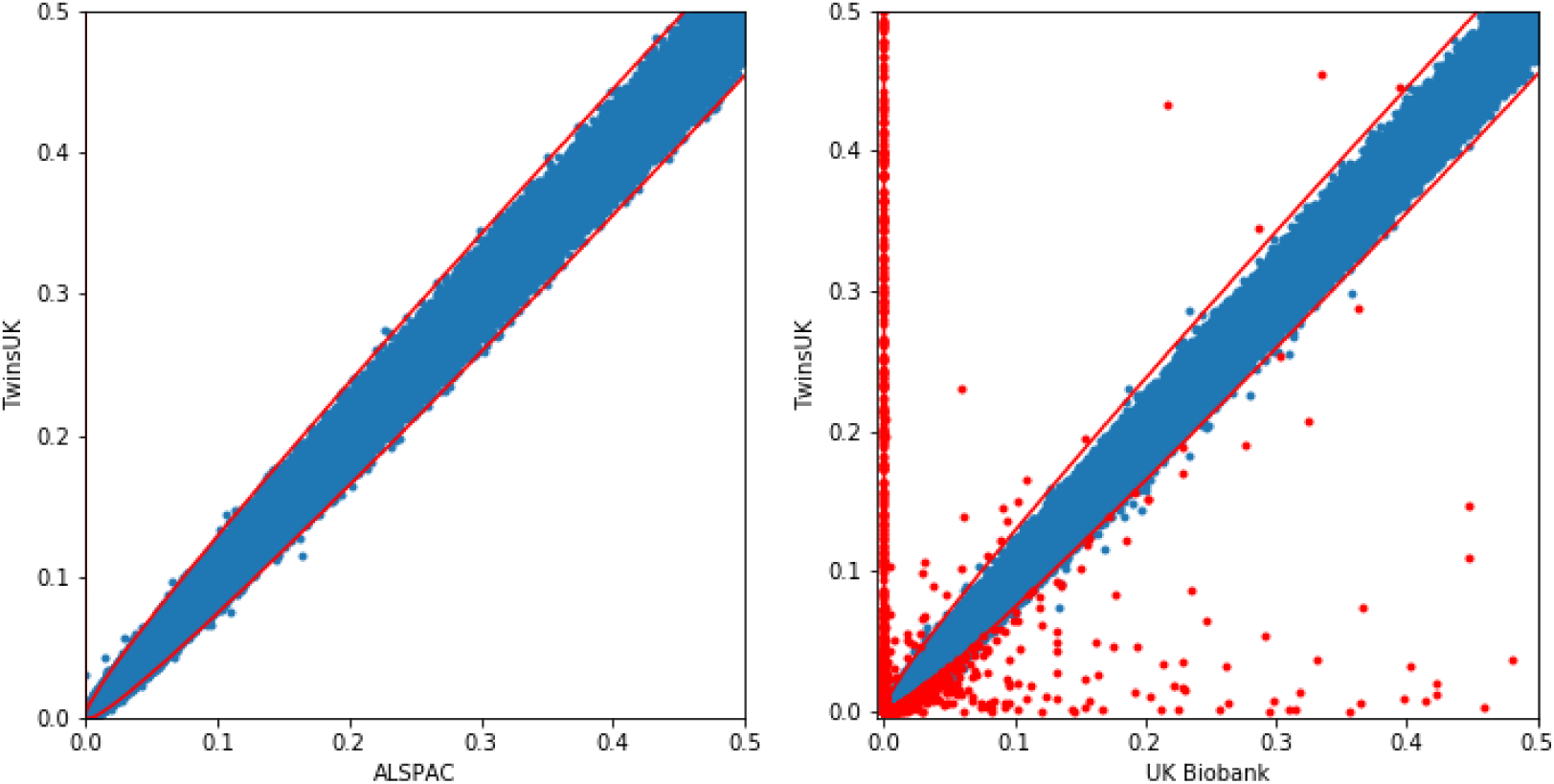
**Left:** The MAF values measured by TwinsUK plotted against those measured by ALSPAC. The red lines shows the threshold at which the binomial distribution has 1-sided p-values of 0.005*/N*_*snp*_. These binomial curves match the actual shape of the distribution fairly closely. **Right:** MAF values of TwinsUK plotted against those measured by the UK Biobank. Many points appear outside of the boundary. The points labeled in red are those SNPs whose UK Biobank MAF value disagrees with both the ALSPAC and TwinsUK MAFs.

We are not aware of any standard pre-processing filtration method that, when calculated from the SNP data alone and without consulting an external database, can reliably distinguish these “out-of-bound” (OOB) SNPs from those that exhibit compatible MAF’s with other studies. For example, similar proportions of both OOB and non-OOB SNPs are filtered out by putting a threshold on the p-value for Hardy Weinberg Equilibrium, so this common preprocessing step, as expected, is not effective in dealing with this problem.

Another method for filtering out potential mismappings is to filter out SNPs that have little or no linkage disequilibrium with their close neighbors. To investigate the effect of this method, we calculated the normalized Mutual Information (MI) between each OOB SNP and all of its neighbors within a distance of 10kb. We used the maximum of these MI values as a measure of local LD; if this value is low, then a SNP is not in LD with any of its neighbors, which may suggest mismapping. Figure 4 shows the distribution of local LD for both the OOB SNPs and an equivalent number of randomly-selected control SNPs. Most OOB SNPs exhibit a similarly strong level of local LD to the randomly selected SNPs, except for a peak in the OOB distribution at MI∼ 10^*–*5^.

**Figure 4:**
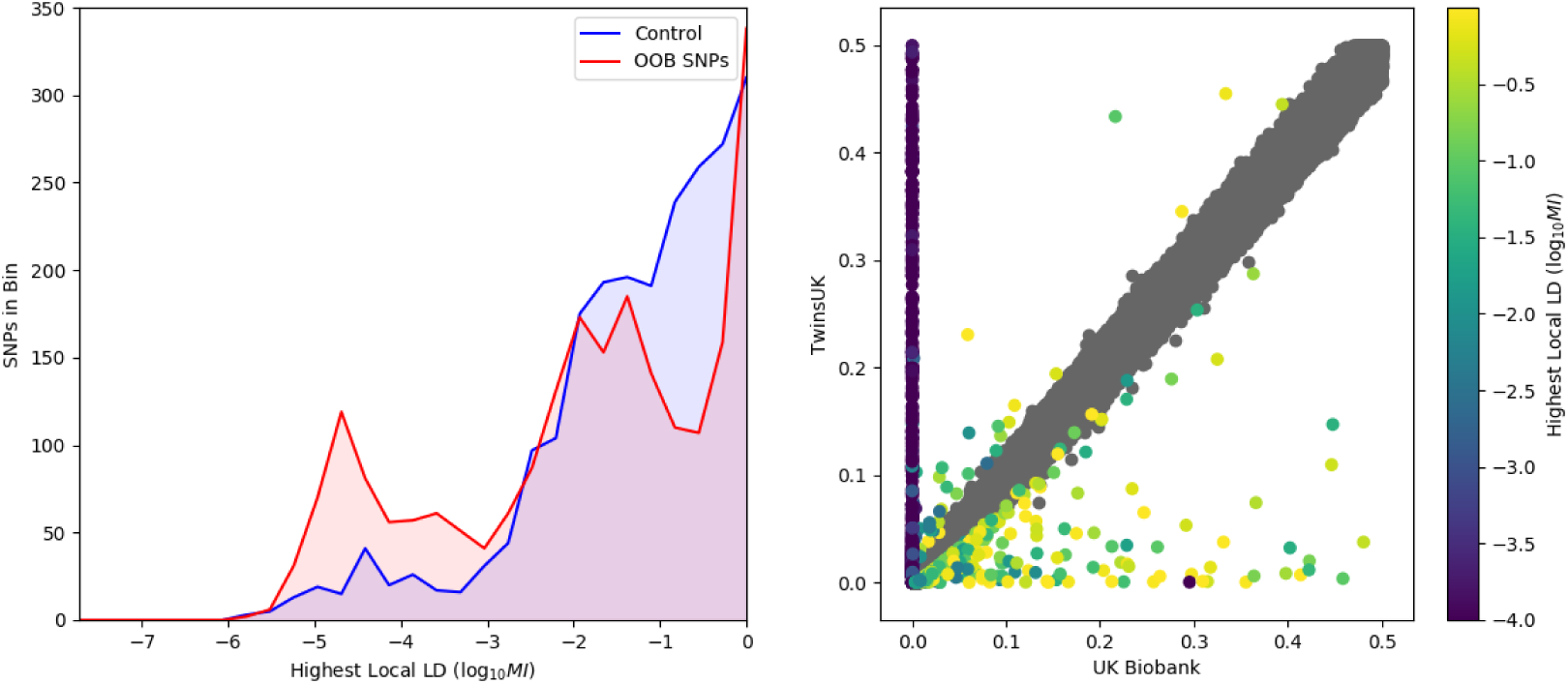
To check for local LD, we calculated the maximum MI between each OOB SNP and their neighbors within 10kb. **Left:** Distributions of maximum local MI for OOB SNPs and random control SNPs. OOB SNPs show similar levels of local LD to the controls, except for a cluster at MI ≈10^*–*5^. **Right:** MAFs of OOB SNPs in both TwinsUK and the UK Biobank (as in Figure 3), color-coded by local LD level. The SNPs with a UK Biobank MAF ≈0 account for almost all of the MI ≈10^*–*5^ SNPs; the SNPs with suspiciously low MAFs also have suspiciously weak local LD. Most other OOB SNPs have unremarkable local LD levels.

In the right panel of Figure 4, we map our measure of local LD onto the TwinsUK vs. UK Biobank MAF plot. This reveals the identity of the SNPs that have what we might call suspiciously weak local LD: the same SNPs have a MAF of ∼0 in the UK Biobank despite having a much larger MAF in both TwinsUK and ALSPAC. This suggests that almost all SNPs that might be filtered out by a threshold on local LD would also be filtered out by a threshold on a minimum MAF value in the dataset. Conversely, the OOB SNPs with a larger MAF in the UK Biobank data tend to have similar levels of local LD to random control SNPs, such that they would not be identified by either filter

We can also consider the UK Biobank’s SNP quality control information (Biobank Resource 1955). The genotyping was performed in several batches, and this resource contains a table indicating whether or not each SNP passed all quality control tests within a particular batch. On average, non-OOB SNPs passed all QC tests (for 99.83% of batches), while OOB SNPs passed all QC tests for 99.5% of batches. Clearly, no strong difference between these groups is detected by these quality control tests.

Based on our observations we propose that comparison with MAFs in external databases is an important quality control step, potentially accounting for difficulties and possible errors that may not otherwise be detected. To this end, we have made these results, including a list of all SNPs identified as OOB, available for download from this link:http://kunertgraf.com/data/biobank.html

## 3 Apparent Interchromosomal Linkage

In the course of our investigations, we found that the SNP rs201947198, for example, violated a usual assumption of genetic analysis: namely that it should be in linkage equilibrium with all SNPs on different chromosomes. To our surprise we found evidence of significant interchromosomal linkage between this SNP and several other markers. To get a comprehensive view of this phenomenon, we calculated the Mutual Information between interchromosomal SNPs across a wide sampling of the database.

Our preliminary analysis took a random subset of 20k individuals and 20% of all SNPs and calculated the MI of every interchromosomal SNP-SNP pair. This resulted in a variety of large, complex networks of interchromosomal linkage, joining sets of SNPs spanning many chromosomes; however, the implicated SNPs correlated with some ethnic phenotypes, leading us to suspect that this initial result was an artifact of the population structure (indeed, the UKB genetic data does encode significant latent population structure [8]). To remove the effects of the population structure, we restricted our full calculation to a strict subset of the population, choosing from the subpopulation meeting the following criteria:

- Self identified as “White British” in the Ethnic Background data field (field 21000)
- Self identified as “Fair” in the Skin Colour data field (field 1717)
- Were not related to other study participants as per the Genetic Relatedness Pairing field (field 22011)
- Were genotyped using the UK Biobank Axiom Array (field 22000)

From the full population, this gave us 222,682 individuals matching all the criteria. From these we extracted a random sample of 10k individuals and calculated the normalized SNP-SNP MI for the full set of all SNPs. SNPs were also filtered out using the following criteria, calculated on the random sample of 10k individuals:

- Minimum minor allele frequency: *f*_*ma*_ ≥ 0.01
- Minimum Hardy-Weinberg p-value: *p*_*hwe*_ ≥ 0.05
- Maximum percentage of missing data: *p*_*miss*_ ≤ 0.05

After filtering out SNPs using these criteria, we then took the remaining SNPs and calculated the MI of every interchromosomal SNP-SNP pair. A randomly shuffled version of this procedure, to gauge a random level, yielded a top MI value of 0.019, and we therefore chose a threshold of 0.03 to score a pair as having “interchromosomal linkage” for the purposes of our analysis. This is an imprecise approach to the statistical significance of these connections, but is mitigated by the sheer strength of many of these associations; the associations of interest to us here are typically an order of magnitude larger than this threshold (see figure 5).

**Figure 5:**
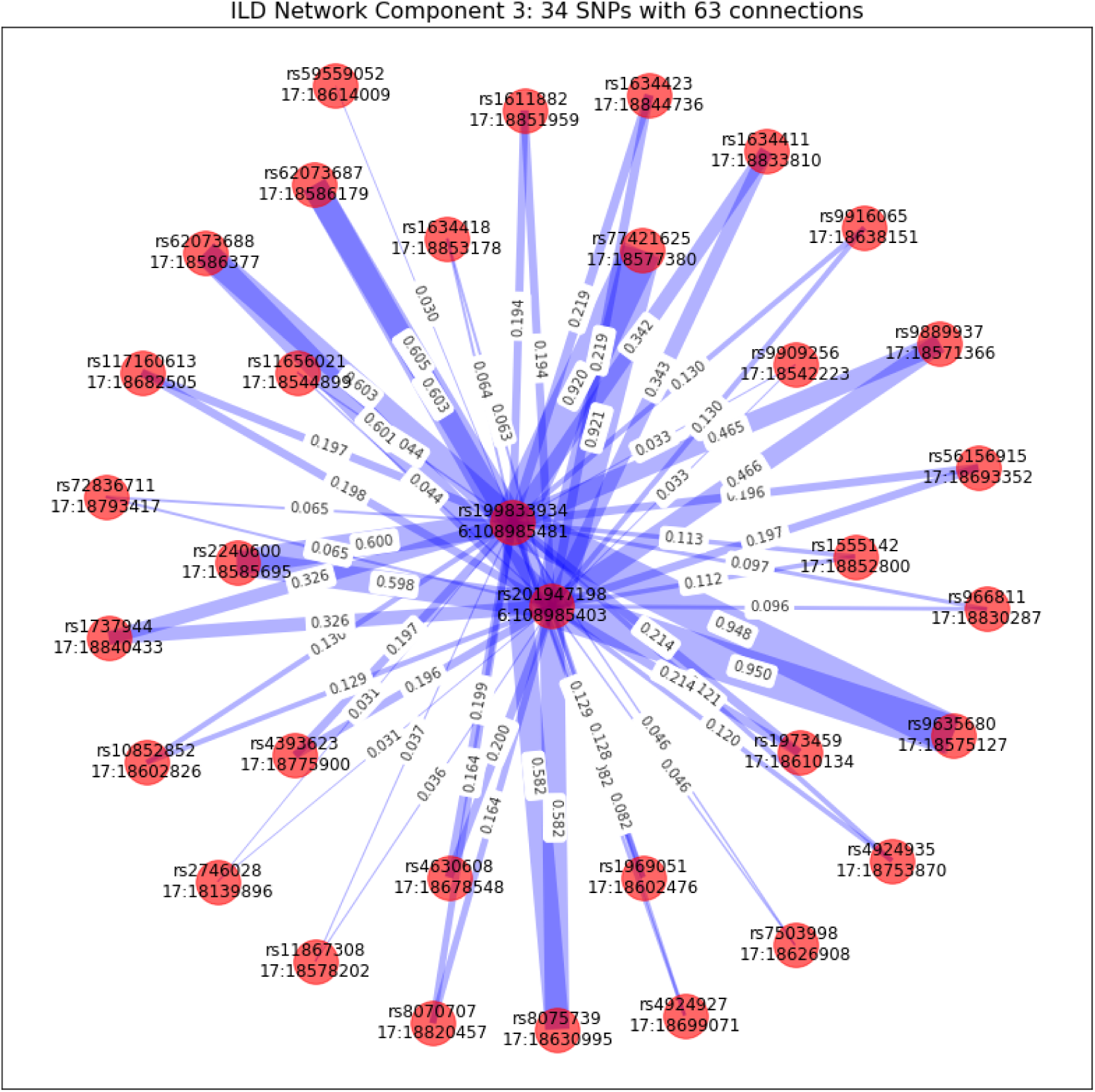
Sample subnetwork of interchromosomal association. A total of 20 disconnected subnetworks were found, all of which share the following basic structure: Each subnetwork includes SNPs from exactly two different chromosomes, and the network structure is always many-to-few (in this case, there are 32 SNPs on chromosome 17 that connect to only 2 SNPs on chromosome 6).

Compared to the preliminary analysis of a random population subset (i.e. networks calculated without filtering out population structure), the network of linkages found here has a fairly simple structure. It consists of 1894 different SNPs that are connected together in 20 independent, disconnected components. A sample of these component subnetworks is shown in Figure 5, and this sample is representative of the simple structure seen in all other components. Specifically, all 20 components consist of SNPs from exactly two different chromosomes, with one chromosome containing many SNPs and the other only a few. For example, the network in Figure 5 has 32 SNPs on chromosome 17 that connect onto only two SNPs on chromosome 6. The question here, of course, is: what could account for this apparent evidence of interchromosomal linkages.

Careful investigation of these SNPs revealed the likely source of the apparent association. For example, in Figure 5 the SNPs on chromosome 6 lie in the gene FOXO3, and the SNPs on chromosome 17 lie in or near the associated, similar gene FOXO3B. On examination of the sequences in these regions there are large segments of sequence that are almost exactly the same (the reverse complement, if we take the strands as presented in the genome builds.) Within the assembly GRCh37.p13, there is a 99.6% reverse-complement match between the 916b sequences at chr6:108985176-108986092 and chr17:18574489-18575405. A portion of this matching sequence, as seen in the NCBI’s Variation Viewer, is shown in Figure 6.

**Figure 6:**
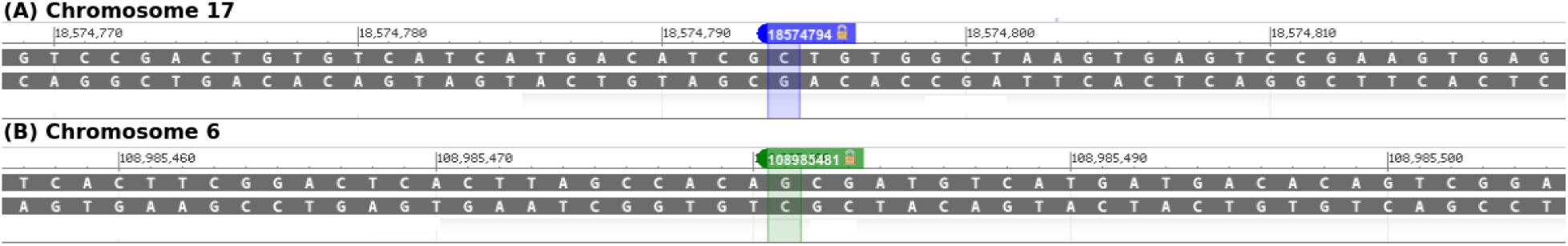
A portion of the sequences inside of FOXO3 and FOXO3B, which are exact reverse complements of each other. There is a 916-base sequence that is 99.6% identical. On Chromosome 6, this sequence contains both the SNPs rs201947198 and rs199833934.

We have examined the “linked” SNPs of Figure 5 carefully and found similar matching subsequences for 19 of the 20 components (and we suspect that the remaining component has a similar underlying sequence pair). Out of the 20 components, 11 consist of SNPs lying on or near previously-identified gene/pseudogene pairs with common sequence (similar to the FOXO3/FOXO3B pair above, though FOXO3B is a functional homolog, not a pseudogene). We did not find annotated gene/pseudogene pairs for the other 8, but sequence scanning (with BLAST) revealed that their SNPs lay in and near common sequences.

### 3.1 Interchromosomal Linkage and MAF mismatches

The subnetwork in Figure 5 includes the SNP rs201947198, discussed previously, which is reported by dbSNP to have a MAF of 0.00002, but has a MAF of 0.375 in the Biobank data. rs201947198 is located at chr6:108985403, which is inside the range of chr6:108985176-108986092 discussed in the previous section. As noted there, this is identically the sequence at chr17:18574489-1857540, in which an identical SNP, rs9635700, occurs at chr17:18574872. Three studies in dbSNP report rs9635700 as having a MAF range of 0.315-0.41, which is consistent with the Biobank’s MAF for rs201947198. Given its previously-reported rarity we suspect that only a negligible portion of the Biobank cohort actually has the minor allele for rs201947198, and that this is instead a mismapping in which the array is actually measuring the minor allele of the identical SNP rs9635700. Therefore the apparent evidence of interchromosomal linkage here is merely the usual intrachromosomal linkage between rs9635700 and its nearby SNPs on chromosome 17, together with a related but spurious signal from another chromosome. Such a mismapping because of the occurrence of common sequences explains both the apparent interchromosomal association and the MAF disagreement between different studies.

We now ask whether all of these interchromosomal mismappings correspond to a MAF mismatch. As shown in Figure 7, this turns out not to be true in general. This figure compares the MAFs measured by the UK Biobank to the most closely agreeing value reported in dbSNP^1^. The left panel of this figure shows the previously-discussed component, Component 3, for which the mismapped SNPs have apparent MAFs that disagree strongly to those reported in dbSNP. In Component 5 (Figure 7, right), however, the mismapped SNP has a very similar MAF to those previously reported. Therefore, these mismappings can not be filtered out by simply filtering out the previously-identified OOB SNPs.

**Figure 7:**
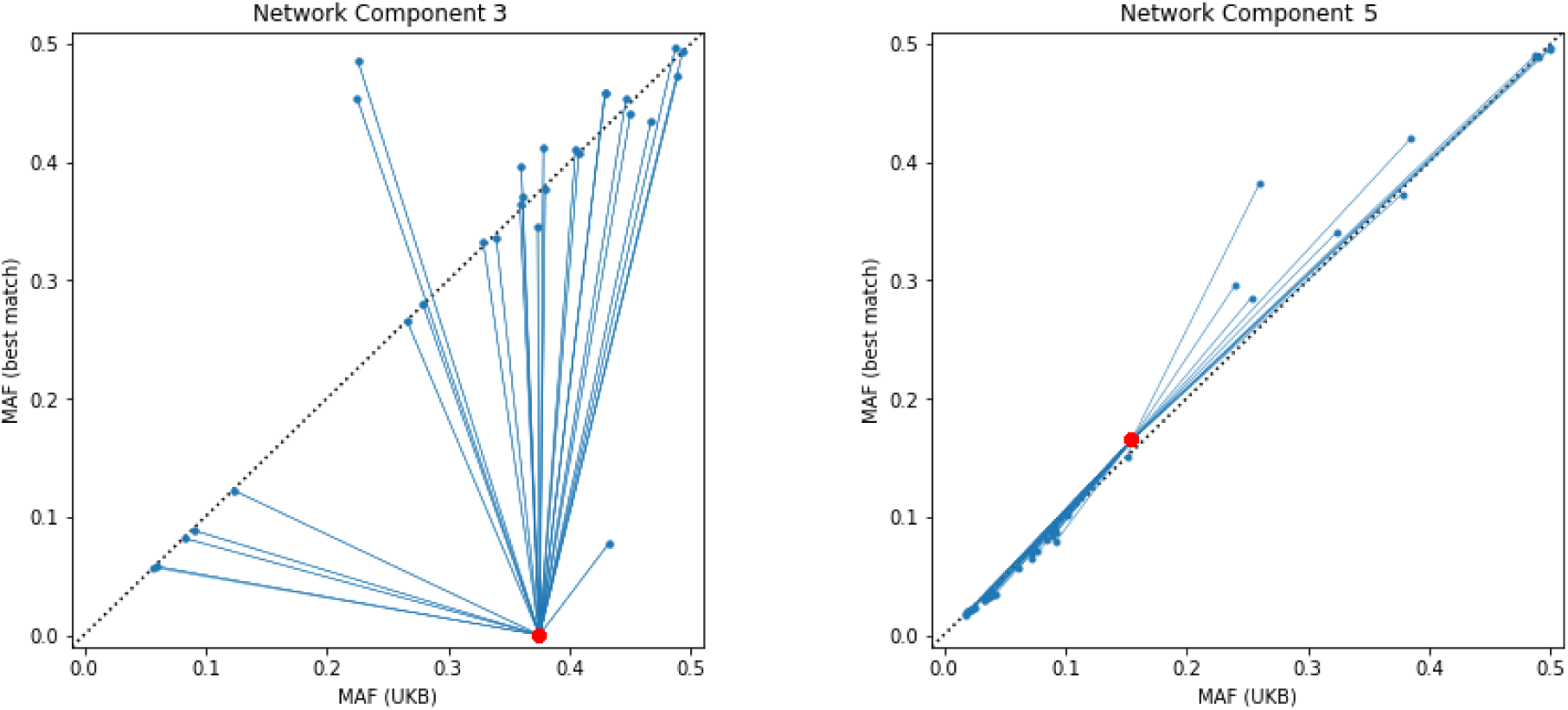
**Left:** the MAFs of the SNPs in the Figure 5 subnetwork, both in the Biobank and in dbSNP. For the vertical axis, we use the MAF value in dbSNP that agrees most closely with the Biobank’s value for that particular SNP. The red dots are the Chromosome 6 SNPs, which (as discussed) vary wildly from dbSNP’s values. **Right:**The same plot for a different component subnetwork, this one linking 12 SNPs on Chromosome 1 to a single SNP on Chromosome 9. The mismapped SNP on Chromosome 9 has a MAF that agrees closely to the one recorded in dbSNP.

## 4 Conclusion

We have identified the following two patterns in the UK Biobank data, implicating SNPs that we propose should be filtered out from analyses, particularly those searching for multigenic interactions:

- The allele frequencies of several SNPs are inconsistent with those measured by two previous studies, ALSPAC and TwinsUK, which were performed on similar populations in the UK
- After accounting for population structure, there remains a set of residual, apparent interchromosomal linkage that appear to be the result of mismappings

There are a number of future directions that may better characterize these phenomena. For example, we do not consider the effects that these SNP mismappings will have upon genomic imputation. Another implication that we have not explored is that our ILD networks appear to effectively detect duplicate genetic sequences within the sample population without requiring full sequencing data; this may be useful in detecting variations in segmental duplications between different populations.

We have generated lists of SNPs implicated by each phenomenon for the purpose of filtering these SNPs out in our pre-processing. For the benefit of researchers who may wish to carefully monitor the SNP data they use and perhaps implement a similar pre-processing step, we have made these lists freely available to download at: http://kunertgraf.com/data/biobank.html.

## 5 Acknowledgements

We would like to acknowledge helpful comments and feedback on earlier versions of this manuscript from Greg Carter, Joe Nadeau, Claudia Fonseca, Rory Collins, and Jonathan Marchini. We would also like to acknowledge help from Elaine Skeffington in editing the manuscript. This research has been conducted using the UK Biobank Resource under Application Number 33492.

## Footnotes

1 For example: the SNP rs71238527 has a MAF=0.216 in the UK Biobank. dbSNP reports values from two different studies, ExAC and GO-ESP, with MAF values of 0.145 and 0.086. We take the closest value, 0.145, as the “best match” from any dbSNP study.

## Notes

http://kunertgraf.com/data/biobank.html

### Competing Interest Statement

The authors have declared no competing interest.

